# Configurational entropy-based screening and selection of hydrophilic polymers using the tripartite split green fluorescent protein

**DOI:** 10.1101/2022.11.07.515508

**Authors:** Shounak Banerjee, Yury Minko, Elizabeth S Anaya, Zachary J Sasiene, Jurgen G Schmidt, Charlie E M Strauss, Geoffrey S Waldo

## Abstract

Measuring the entropic properties of polymers such as proteins is critical to accurate prediction of their functional properties. However, the measurement of configurational entropy is possible only by low throughput techniques such as calorimetry, NMR and CD spectroscopy. Moreover, to our knowledge no system exists that allows molecular selection/enrichment based on the molecules configurational entropy. We tested the ability of the scalable tripartite GFP system to offer fine resolution of differences in configurational entropy in molecules and to isolate molecules based on their configurational entropy. The system was able to both finely resolve molecules with different configurational entropies, as well as capture them for isolation. We were able to tune the sensitivity of the system by using different mutations of the protein components. Lastly, we were able to apply the system to polypeptoid molecules and posit that the system may be applied to any other hydrophilic polymer of up to 10^3 repeating units.

## Introduction

Proteins are the functional units of life. They are also commonly used industrially for stereoselective catalysis^1,2^. Most proteins are heteropolymers of amino acids linked by peptide bonds. There are twenty amino acids commonly found in all organisms. They all share a common backbone of a carbon atom flanked by an amide and carboxyl group. This carbon atom can have different functional groups, which give each of the twenty amino acids their unique identity and physicochemical properties. The directional arrangement of the amino acids in a protein, also known as the protein’s primary sequence, dictates its structure and functions^3^.These functions, like any thermodynamic process, are achieved by balancing enthalpy and entropy. This balance is accomplished by altering the conformational ensemble of the protein to gain or lose intrachain interactions in favor of interactions with the binding partner, which may be a solute, solvent or other polymeric molecule^4–6^. While proteins are excellent examples of molecular recognition agents, they have drawbacks such as susceptibility to biological degradation, limitations on amino acid choices and therefore explored structure and function space. To address these drawbacks and develop more versatile molecular recognition and catalysis agents, there is growing interest in proteomimetic polymers^7–12^. In order to develop predictive models for proteomimetic behavior, the knowledge and understanding of their thermodynamics, namely enthalpy and entropy changes for structure modulation is essential. Thus far, enthalpy and entropy changes in molecules have been measured using calorimetry^13–15^, circular dichroism^16–18^ and nuclear magnetic resonance (NMR) spectroscopy^19–21^. These methods do not allow high throughput measurements nor simultaneous selection or enrichment of molecules in a library based on their configurational entropy. They also measure bulk properties and cannot be applied to single molecule studies or isolations. Though high-throughput methods for measurement and selection based on configurational entropy exist, they can only be applied to proteins and are low-resolution, distinguishing between structured and unstructured polypeptides^22^. Using current high-throughput configurational entropy-based selections, it is not possible to obtain a continuum of proteins with various degrees of structure. Techniques such as phage display^23,24^, yeast surface display^25^ and ribosome display^26^, that are commonly used to select molecular recognition agents such as antibodies are also not suitable for configurational entropy based selection as they implicitly rely on cellular machinery that degrades misfolded or otherwise unstructured peptides. The lack of high throughput screening and selection methods based on configurational entropy precludes accurate predictions of the thermodynamic properties of protein interactions with their targets. It is even more problematic for the design of proteomimetics, where it is desirable to have a screening and selection method that is functional across a wide gamut of configurational entropies, to capture and sample a diverse set of molecules.

In prior work^27^, we have shown that the tripartite GFP system can distinguish between two configurations of a sulfite reductase GFP10-GFP11 fusion, producing a higher signal when the GFP10 and GFP11 are fused as a continuous hairpin than when the GFP10 and GFP11 are at the N and C termini of the sulfite reductase protein. Ostensibly, this is due to the higher configurational entropy of the GFP10 and GFP11 when fused to the termini of sulfite reductase. Therefore, we reasoned that the tripartite GFP system may be able to measure and distinguish between polymers with more finely varying amounts of configurational entropy. To test this hypothesis, we performed two experiments. First, we inserted tetrapeptide blocks of three glycines, followed by a serine, between GFP10 and GFP11. Since glycine and serine rich linkers are known to be disordered, we expected that configurational entropy would substantially increase with the addition of more tetrapeptide blocks. In the second experiment, we genetically fused GFP10 and GFP11 to the termini of the repeat in toxin (RTX) domain, previously characterized by Szilvay et al^28^ and titrated calcium chloride which binds to and induces folding of the RTX domain. In both cases we expected the tripartite GFP to show decreasing rate of complementation with increasing configurational entropy.

Here we show that the tripartite split-GFP system behaves as expected and is therefore capable of measuring fine changes in the configurational entropy of a hydrophilic polymer as well as selecting the polymer from a library of polymers. Figure 1 depicts a model for the system’s operation. Here the GFP10 and GFP11 may be synthesized chemically and conjugated to any hydrophilic polymer of interest or a library of polymers. Notably, such a library can be barcoded, allowing determination of the molecular identities for polymers of interest. We also show that the sensitivity of the system is tunable, allowing configurational entropy measurements across a wide gamut of configurational entropies and demonstrate the utility of this system with polypeptides and polypeptoids. Currently, using the system with polypeptoids and other abiotic polymers requires the GFP10 and GFP11 to be chemically synthesized, either in-line with or to be conjugated to the polymer. The synthesis yields are sensitive to the presence of sulfur containing amino acids, namely methionine and cysteine. GFP11 has a methionine as its fourth residue. We therefore asked whether this methionine could be substituted with its sulfur-less isostere, norleucine for improving the synthesis yields. Prior work in the Waldo lab has found that most mutations of this methionine are highly deleterious for GFP function. Therefore, we were surprised to find retention of significant fluorescent function with the non-canonical amino acid substitution. We then demonstrated the ability of the tripartite GFP system, with the norleucine substitution, to distinguish between sarcosine oligomers of different lengths.

## Results

### Initial rate of *in vitro* fluorescence gain decreases with addition of entropy blocks

We found that using different combinations of GFP1-9 and the GFP11 (Table S3) produced dramatic changes in the response to variation of chain length. For complementation reactions with 1-9 OPT Fast, the complementation rate was found to decrease approximately log-linearly (R^2^>0.96) with the number of added entropy blocks (Fig. 2).

**Figure 1:**
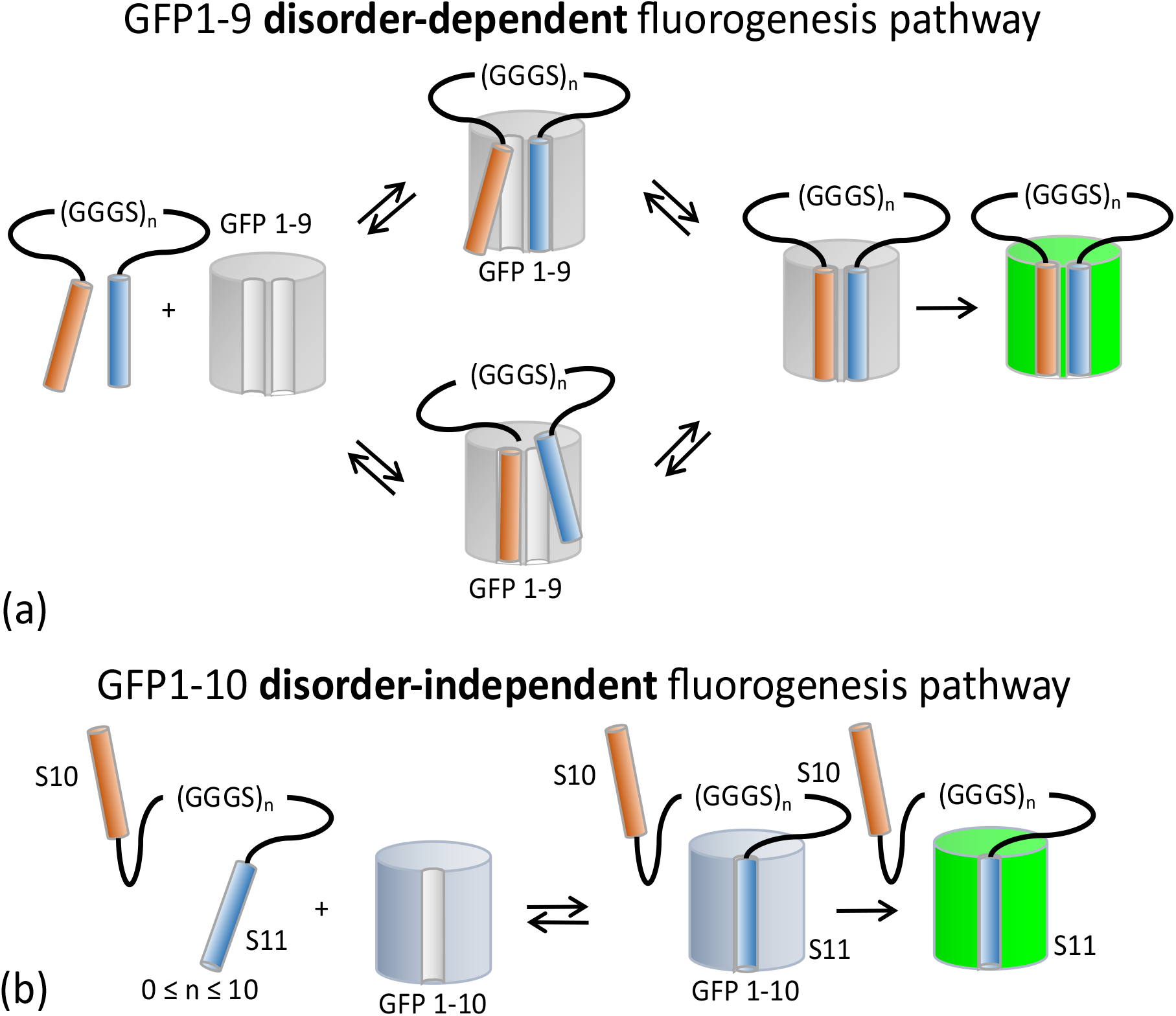
Mechanism of (a) disorder-dependent complementation of analyte end-labeled with S10 and S11, with GFP1-9 that produces fluorescent signal. This is contrasted with (b) disorder-independent complementation of the same analyte with GFP1-10, which also produces a fluorescent signal.

**Fig. 2:**
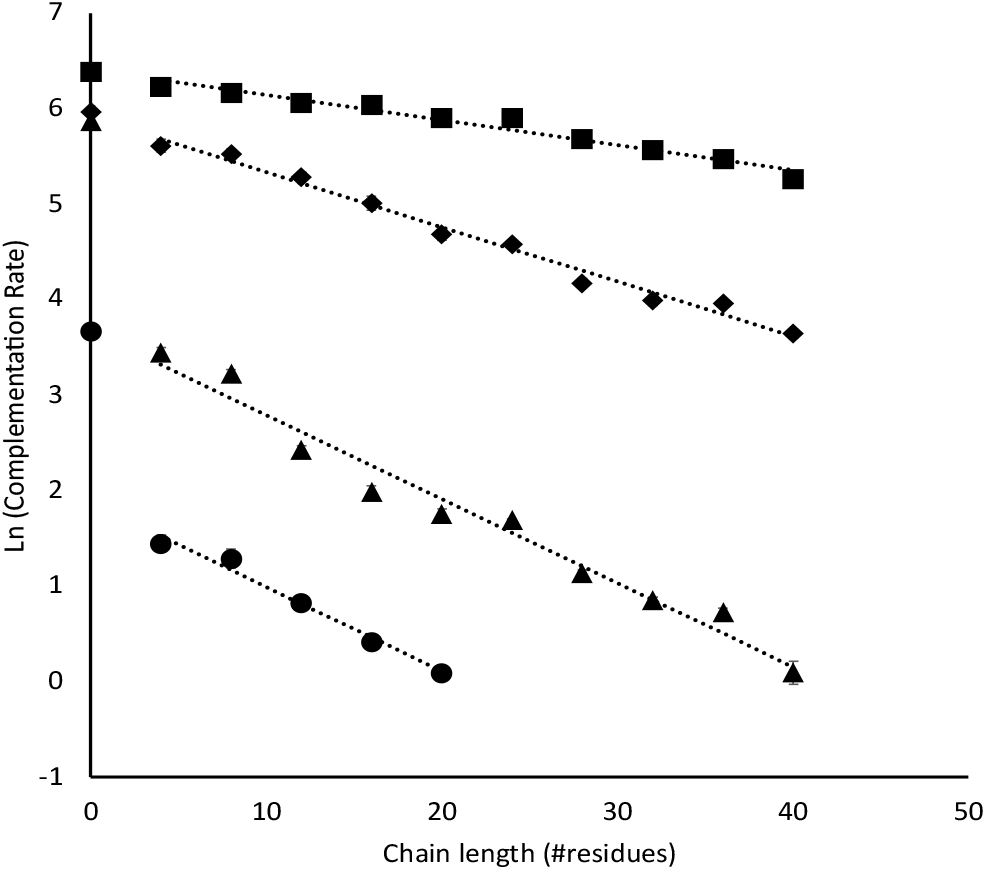
Log -linear plots of the complementation rate (calculated between 30 and 50 min, post-initiation of complementation), as a function of the logarithm of chain length, for different combinations of GFP1-9 and GFP11. Fast 1-9 and Fast 10-11 (squares), Fast 1-9 and Slow 10-11 (diamond), Slow 1-9 and Fast 10-11 (triangles) and Slow 1-9 and Slow 10-11 (circles). Dotted lines represent linear fits to each data series.

For complementation reactions with 1-9 M1 slow the 10-11 hairpin (0 block) constructs, displayed a significantly higher complementation rate than predicted by calculating the intercept of extrapolations from log-linear fits to the data from the rest of the constructs (1 block – 10 block). The results were consistent with our previous study^27^ that measured complementation kinetics for 1-9 OPT Fast and 1-9 M1 Slow, with a loop insertion of the GFP10-11 hairpin sequence into sfCherry^29^ and with SR that was end-labeled with GFP10 and GFP11^27^. These previous experiments showed that for 1-9 OPT Fast, complementation rates for the two targets were similar, while 1-9 M1 Slow showed a significant decline in complementation rate for the end-labeled SR, compared to the sfCherry with loop insertion.

### Effect of carrier protein on complementation rate

SR was used to mitigate the potential problem of non-expression of the smallest constructs, acting as an expression driver. To find out whether the driver protein affects complementation, the SR coding sequence was replaced by that of SUMO. The latter fusion proteins were digested by the SENP protease. Cleavage of SUMO from the entropy ladder fragments were confirmed by SDS-PAGE (Fig. S4) and complementation reactions were set up. Undigested SUMO-entropy ladder proteins were used as controls. No significant difference was found in the complementation rates between the two types of constructs (Fig. S2). The N50 values calculated using equations 1 and 2 for the four polypeptide systems are provided in Table 3.

### *In vitro* behavior is mirrored *in vivo*

Next, we sought to determine whether the system could be deployed *in vivo*. To more accurately mimic the *in vitro* experiments, we used sequential induction^30^. Fluorescence was undetectable at the onset of induction for the 1-9 or 1-10 proteins.

We found that akin to the *in vitro* experiment results, fluorescence intensity declined with increasing chain length of the entropy ladder (Fig. 3).

**Fig. 3:**
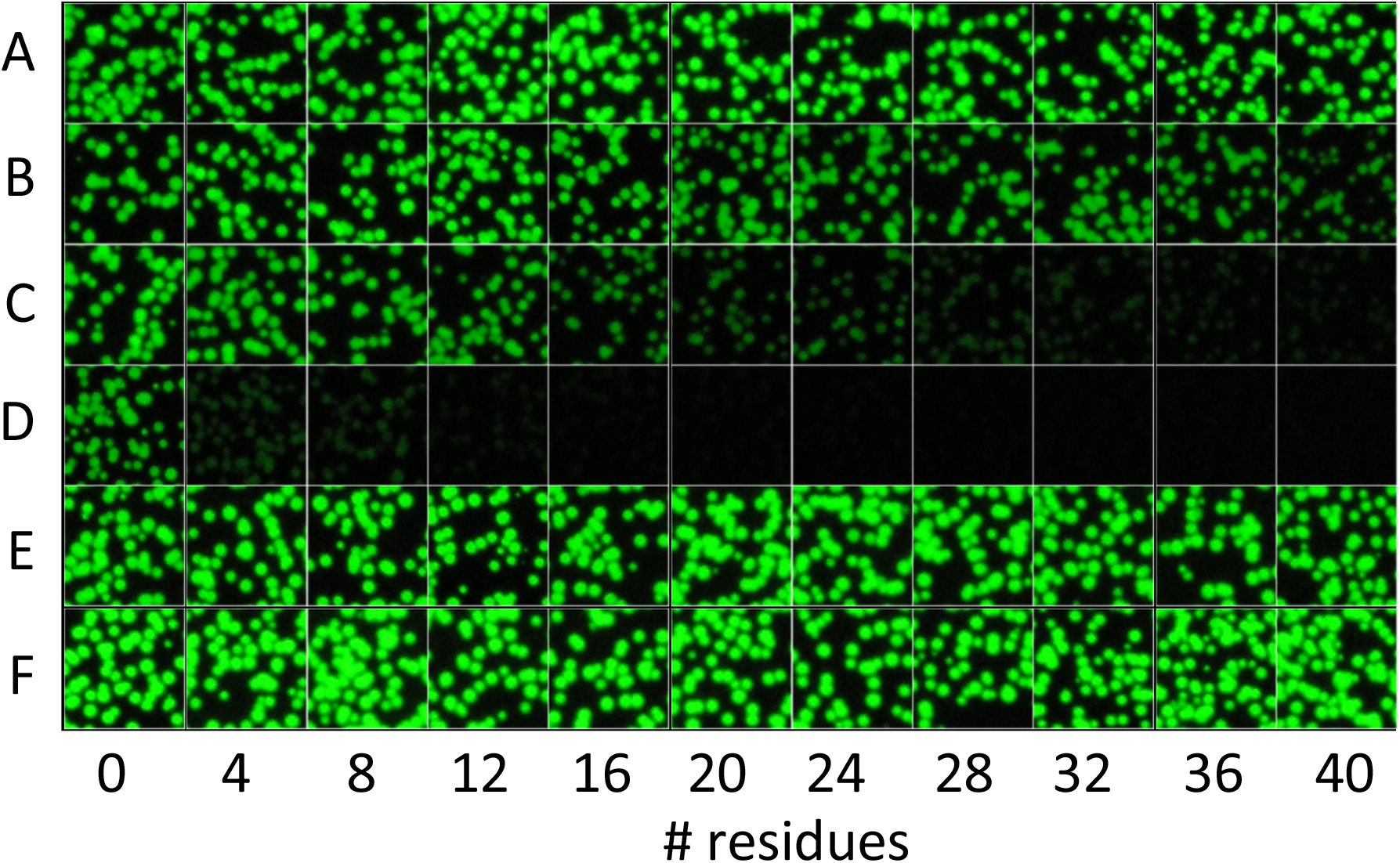
Fluorescent bacterial colonies after sequential induction of entropy ladder (2 h) and GFP 1-9 or GFP1-10 (2 h). Different rows shown are for cells expressing GFP1-9 or GFP1-10 variants and variants GFP11 at the end of the entropy ladders. (A) 1-9 OPT Fast and 11 M4 Fast, (B) 1-9 OPT Fast and 11 M3 Slow, (C) 1-9 M1 Slow and 11 M4 Fast, (D) 1-9 M1 Slow and 11 M3 Slow, (E) 1-10 and 11 M3 Slow and (F) 1-10 and M4 fast

The trends in sensitivity were also preserved, with the combination of 1-9 OPT Fast and 11 M4 Fast showing no significant decrease in fluorescence (and therefore complementation rate/efficiency) over the range of chain lengths examined; whereas the combination of 1-9 M1 Slow and 11 M3 Slow was the most sensitive, with fluorescence rapidly falling off with the addition of one entropy ladder block.

Complementation was allowed to continue up to 24 h. Then we imaged the colonies again, using the procedure described above. These images, of equally fluorescent colonies, are shown in Fig. S5.

### Low entropy constructs out-compete high entropy constructs in competitive binding assays

Competitive binding experiments were set up with the mixtures outlined in Table 4. The experimental design and results are presented in Fig. 4.

**Fig. 4:**
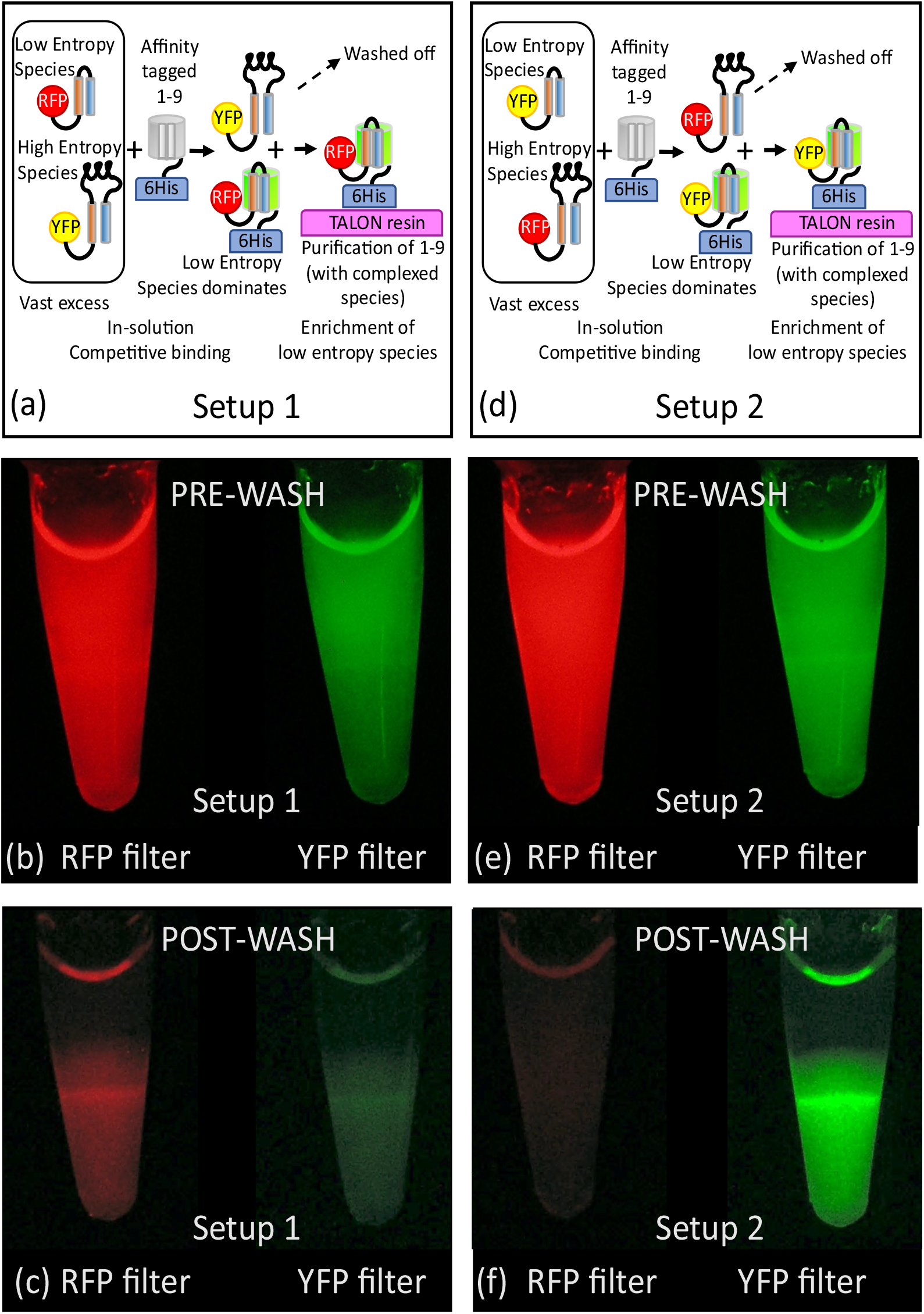
Enrichment of low-entropy polymers. In Setup 1 (a), a red fluorescent protein (sfCherry) is attached to the GFP10-11 hairpin (low entropy) construct while a yellow fluorescent protein (YFP-OPT2) is attached to the 10 block entropy ladder construct (high entropy). (b) Fluorescence images captured using the RFP filter and the YFP filter (Table 2), after loading of constructs pre-complemented with polyhistidine tagged GFP1-9 M1 and after washing off unbound material (c). In Setup 2 (d, e, f), the same setup and procedures as Setup 1 but sfCherry is attached to the high entropy construct and YFP-OPT2 is attached to the low entropy construct.

Fluorescence images after loading TALON resin for Setups 1 and 2 (Fig. 4c and 4e) show that approximately equal amounts of each labeled species were present. Only red fluorescence was observed on the resin when sfCherry was attached to the 0 block entropy ladder (Fig. 4c) and only yellow fluorescence was observed with YFP-OPT2 was attached to the 0 block entropy ladder (Fig. 4f).

### Detection of calcium induced folding of repeat-in-toxin domain

To test the ability of the system to detect partial ordering in proteins of equal chain length, we used the repeat in toxin (RTX) domain V of B. pertussis adenylate cyclase A (cya A). Like Szilvay et al,^28^ we refer to this domain as RTX-BR(L). The SR-10 block entropy ladder was used as a non-responsive control.

The *in vitro* complementation assays suggested that the combination of 1-9 OPT Fast and 11 M4 Fast would be able to provide a signal even if the RTX domains were completely disordered. We therefore used this combination of GFP1-9 and GFP11 for this experiment. RTX-BR (L) showed significant improvements in complementation rates with increasing calcium chloride concentrations, compared to the non-responsive SR—10 block entropy ladder construct. The latter showed an increase in complementation rate at calcium chloride concentrations above 1 mM (Fig. S2). Fitting of the titration data (Fig 5) to a Hill equation for cooperative binding yielded a dissociation constant (KD) value of 0.202 mM and a Hill coefficient of 4.2. Our calculated dissociation constant value was in quantitative agreement with that calculated by Szilvay et al.

**Fig. 5:**
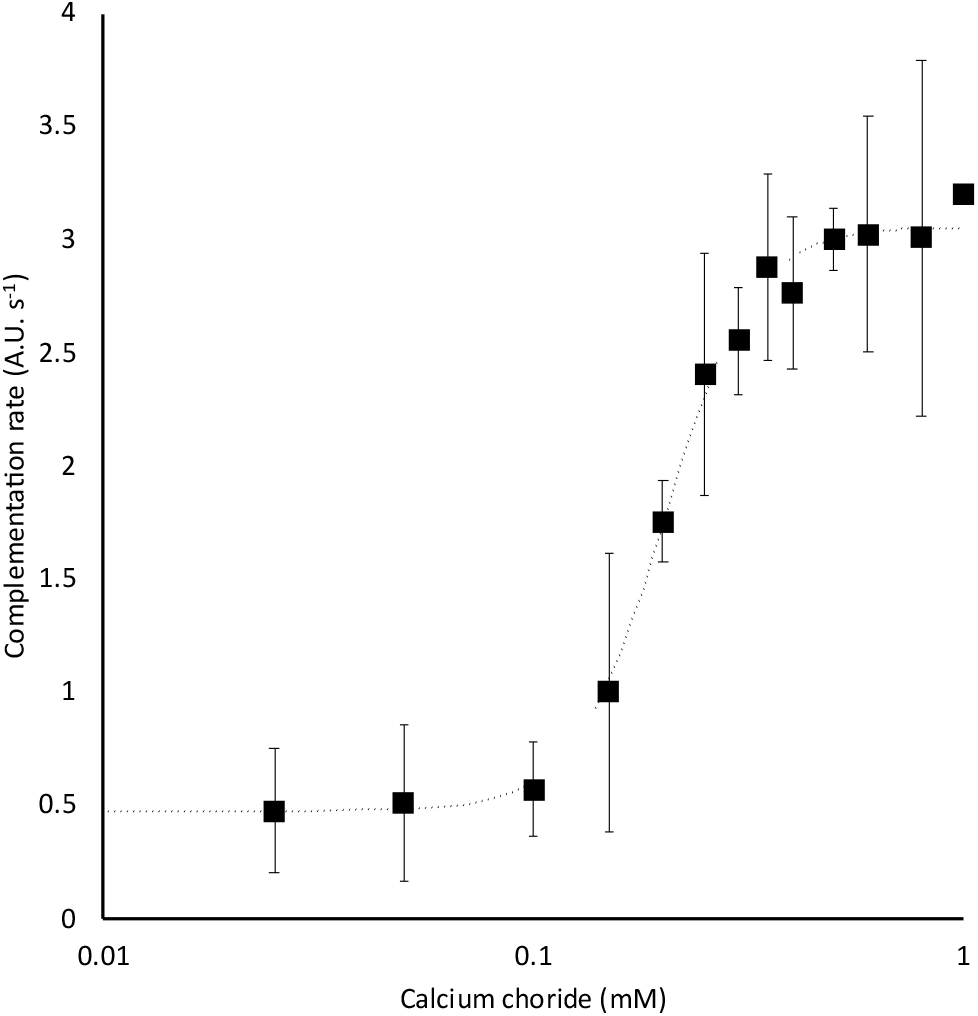
Titration of calcium against RTX-BR(L) measured with GFP1-9 OPT Fast and GFP11 M4 Fast. The fit of a Hill model, to the data is shown as a dotted black line.

The contrast between complementation rates with and without 6 mM calcium chloride was greater after omission of glycerol from the buffer. (Fig. S1)

### End labeling with synthetic GFP10 and GFP11 strands enables characterization of peptoid oligomers

We synthesized GFP10 and GFP11 using solid phase methods. To improve the yields of GFP11, M218 was replaced with norleucine, thus avoiding oxidation commonly encountered on methionine residues in solid phase syntheses. M218 is a buried residue and is highly conserved. We found that substitution of M218 with any other residue decreased complementation efficiency (Waldo, unpublished data). However, substitution with norleucine produced faster binding kinetics than GFP11 M4, when titrations were performed against GFP1-10, as shown in Fig. S6. We also determined that mixed with GFP1-9 and GFP10 in a tripartite mixture, this synthetic GFP11 did not produce a fluorescent complex. Encouraged by these results, we conjugated this M218Nle variant of GFP11M4 and GFP10 to the ends of oligo-sarcosine peptoids. *In vitro* complementation assay results (Fig. 6) showed that complementation rate decreased with increasing chain length, as observed for the peptide entropy ladders. The results were qualitatively consistent with the findings of Sisido and coworkers.^31^ The sequences for the peptomers (peptide-peptoid copolymers) is provided in Table S2.

## Discussion

### System dynamics and scope of use

Loop closure rates in molecules similar to the entropy ladder constructs are known to decrease as power-law functions of the number of residues^32–34^.In our system, loop closure ostensibly results in the juxtaposition of GFP10 and GFP11, enabling binding to GFP1-9, initiating irreversible fluorophore maturation and resultant fluorescence.

The fluorescence evolution trajectories are similar to the ones seen in previous work in that they asymptote to a maximal fluorescence intensity, which is proportional to the amount of analyte added (data in ESM). This fluorescence does not decrease when residual analyte is removed, with the complex immobilized to metal affinity resins (data not shown). This suggests that the 1-9:10-analyte-11 complex is stable and unlikely to dissociate (Waldo, unpublished data). The system therefore acts as a trap for closed-loop conformers with k-2∼0. This would lead to depletion of free entropy ladder molecules, which is different from other measurements where no external constraints are imposed on the analytes and where an equilibrium is established between open and closed loops. We therefore expected the complementation rate to be proportional to the rate of loop closure and decrease monotonically but not as a power law function of chain length. A log-linear decline in complementation rate with chain length was observed for experiments with 1-9 OPT Fast. For 1-9 M1 Slow, the complementation rate observed for 0 block (native GFP10-GFP11 hairpin) constructs was much higher than predicted by the linear fits of the data. This can be due to the persistence of a structured and binding competent state, driven by inter-residue interactions or inter-chain interactions between the 1-9 M1 and the loop connecting strands 10 and 11. These interactions are broken by adding a single entropy block, which the 1-9 M1 slow is sensitive to. Flattening at longer chain lengths is also observed for complementation reactions with 1-9 M1 Slow and can be attributed to the absence of detectable levels of assembled (and therefore fluorescent) 1-9: Entropy ladder complexes.

The system does not provide mechanistic insight on the relative magnitudes of k1 and k-1. However, this is not the intended use of this system. This system can instead be used to finely distinguish between polymers with different configurational entropies and affinity-purify polymers with desired configurational entropy from a library of polymers.

We used repeats of the tetrapeptide -G-G-G-S-in our experiments to minimize the propensity to form stable structured states. This polypeptide composition is different from the -G-S-sequences that are well studied. It is known that varying the amount of glycine in a glycine-serine linker affects loop closure rates.^33,35^ For this reason and the trapping of closed loops described above, we cannot directly compare our results with the literature. Instead, we intend for this system to be used as a comparative and selective tool in screening libraries of designed polymers of the same chain length.

It is important to also use the same molar concentrations of each polymer in a test set since concentration is a primary determinant of complexation rate. We recommend using higher amounts of analyte for longer chains as long as the GFP1-9 and GFP1-10 fragments are in 8-fold molar excess. This may be construed as higher concentration, if the path length is fixed, as in a cuvette. If the path length can be varied, higher amounts can also be achieved by using larger volumes. This latter approach may be useful when increasing the concentration is not possible or desirable.

Since the system cannot discriminate between intra and intermolecular interactions, we recommend an additional round of testing for enriched members from a library screen. It is possible for two disordered chains to associate without prior structure formation^36^. We expect non-linear concentration dependent and titration responses to result from intermolecular interactions. The tripartite GFP system can of course, be used in its classic form, with GFP10 and GFP11 on different molecules, to easily detect intermolecular association.

### Tuning entropy sensitivity

In our mechanistic model, k2 is a variable dependent on the choice of GFP1-9 and GFP11 variant and ostensibly GFP10. This provides an opportunity to tune the sensitivity of the system to rates of loop closure. Since different GFP11 mutants show varying responses to the same GFP1-9 mutant it is possible that that k1/-1 is affected by the choices of GFP10 and GFP11 mutants.

When 1-9 OPT Fast is used with 11 M4 Fast, the decrease in fluorescence as a function of chain length is small. This is in agreement with previous findings^27^, where interacting protein pairs were found to produce uniform and robust complementation, only requiring GFP10 and 11 to be colocalized. Linker length was not found to be important except for when protein termini were buried and a long linker was required to make the GFP10 and GFP11 accessible.

When 1-9 M1 Slow is used with 11 M4 Fast, the fluorescence falls off much more rapidly, offering more stringent selection for low-entropy molecules. This increased apparent dependence could be due to the lower affinity of 1-9 M1 Slow for the GFP10-11 M4 beta-strand. Structural inspection of superfolder GFP (PDB: 2B3P) and 1-9 OPT Fast in complex with sfCherry via a GFP10-11 loop insertion (PDB: 4KF5) reveals that a lysine at position 149 on GFP1-9 OPT packs much better against Y200 on GFP10-11 than an asparagine that is unchanged in 1-9 M1 Slow.

Since the GFP1-9 is added in vast molar excess, we assume that folding of GFP1-9 is not limiting and that even though 1-9 M1 slow is less well-folded than 1-9 OPT Fast, a sufficient excess of the former is still available to bind to the target. For the constrained hairpin, 1-9 OPT Fast and 1-9 M1 Slow show different rates of fluorescence gain. This gap must be either maintained or be larger for the unconstrained hairpin unless in the case of constrained hairpin, trapped states of M1 contribute to reduction in complementation rate. If this is true, then we expect the complementation rate for the unconstrained hairpin to be lower than for the constrained hairpin.

In summary, we speculate that in equilibrium, the entropy ladders all sample different concentrations of the loop-closed state. The concentration of the loop-closed state is dependent on both, the configurational entropy of the polymer as well as the choice of GFP10 and GFP11, that the consistent non-linear, monotonic decrease in complementation rate using only different variants of GFP11, can be attributed to. These loop-closed states are bound with different affinities by the different GFP1-9 variants. We speculatively attribute deviations from log-linearity, observed for GFP1-9 M1 Slow, to the possibility that the concentrations of loop-closed states are significantly lower than the dissociation constant (KD) of GFP1-9 M1 Slow and the target. The observation that the lower baseline for GFP1-9 M1 Slow complementation approaches zero, supports this attribution.

### Entropy dependent enrichment

We mixed roughly equimolar amounts of the 0 block (hairpin) and 10 block entropy ladders, so that the only discriminant in ability to bind GFP1-9 protein would be configurational entropy. We use the entropy ladders’ respective fluorescent protein labels for quantification. We recommend preparing a standard curve with the fluorescent protein of interest for this purpose. To verify whether binding is affected by the choice of label, we swapped the red and yellow fluorescent protein labels and the results suggest that binding is independent of the label. In our mechanistic model, k-2 is empirically known to be negligible.This means that the probability of exchange, of a bound low-entropy analyte, with a high entropy is infinitesimally small and given enough time, all entropy ladders can bind to GFP1-9. Therefore, the labeled entropy ladders were added in vast excess of the GFP1-9. We also allowed a relatively short amount time for complementation and then washed off excess protein after loading onto TALON resin. The timings of these steps may be tuned for optimal contrast or enrichment profile. We anticipate that in use cases for enrichment of ordered members of a labeled/barcoded library, high concentrations of highly disordered members in the library can reduce the yield of ordered members, which may be at relatively much lower concentrations. We therefore recommend either representing members equally or using unlabelled/unbarcoded “blocking” constructs with known entropy, to improve enrichment signals.

Detection of disorder-order transitions in RTX domains. Our estimates of RTX-BR(L)’s dissociation constant and cooperativity for calcium binding are in quantitative agreement with that calculated by Szilvay et al. We also noticed that the complementation rates were enhanced in increasing ionic strength (>1 mM calcium chloride) for the non-calcium-responsive SR-10 block entropy ladder construct. However, in the TNG buffer, addition of calcium appeared to have no effect on the complementation of entropy blocks with GFP1-9 (data available upon request). This highlights the importance of appropriately calibrating the system before use in a buffer system other than TNG. We did not test the deactivated RTX-BR (L) mutant in this study but it would be interesting to determine if the difference in its behavior relative to RTX-BR(L) can be captured by our system. Ostensibly, the system can be used to evolve proteins like RTX-BR(L) to tune affinity for metals and small molecules.

## Materials and Methods

### Design and construction of entropy ladders

Coding sequences for GFP10 variant M2, tetrapeptide repeats (GGGS; referred to as disorder-blocks) numbering 0 (native GFP10-11 hairpin) to 10 and either GFP11 variant M3 or M4 were synthesized by overlap extension PCR using the Phusion High Fidelity Polymerase (New England Biolabs, USA). For this work, we shall refer to variants of GFP1-9 and GFP11 by their complementation phenotype (Fast/Slow). Table 1 shows the previously used construct nomenclature and its mapping to the nomenclature used in this study.

**Table 1:** Mapping of names used in prior work to those used in this study

| Name in previous work | Name in this study |
| --- | --- |
| GFP1-9 OPT<sup>27</sup> | 1-9 OPT Fast |
| GFP1-9 M1<sup>27</sup> | 1-9 M1 Slow |
| S11 M3<sup>37</sup> | 11 M3 Slow |
| S11 M4<sup>27</sup> | 11 M4 Fast |
| GFP1-10 D7 OPT<sup>37</sup> | GFP1-10 |

The final PCR products were genetically fused to the C-terminus of the *Pyrobaculum aerophilum* sulfite reductase dissimilatory subunit gamma (SR)^37^ by overlap extension PCR. Links to their coding sequences and resulting polypeptide sequences are provided in Table S1. For a subset of these sequences, SR was replaced with the *Saccharomyces cerevisiae* small ubiquitin-like modifier (SUMO)^38^.

### Cloning and selection

For *in vivo* studies, the genetic constructs outlined above were cloned using 5’ NcoI and 3’ KpnI restriction sites, into the pTET ColE1 vector.^27,30^ Briefly, each insert was double-digested using the NcoI-HF and KpnI-HF restriction enzymes (New England Biolabs, USA) using previously described protocols^37^. The vector was digested with the same enzymes and Calf Intestinal Phosphatase (New England Biolabs, USA) to dephosphorylate 5’ ends.Digested inserts were gel extracted using the QiaQuick gel extraction kit (Qiagen, Germany). Digested and 5’ dephosphorylated vector was purified using the QiaQuick PCR Cleanup kit (Qiagen, Germany). Both inserts and vector were eluted and stored in 10 mM Tris-Cl, pH 8.5 until later use. Ligation was performed with a 3.5:1 v:v ratio of vector to insert, using Rapid Ligation buffer (Invitrogen, USA) and T4 DNA ligase (New England Biolabs, USA). Ligation products were used to transform BL21DE3 Gold (Agilent, USA) cell lines carrying 1-9 OPT fast, 1-9 M1 slow or 1-10 on a pET p15A expression vector^27,30,39^. Transformants were screened by sequential induction^30^ and colonies representing the dominant phenotype were picked and stored for subsequent use, in 25 % glycerol stocks.

For *in vitro* studies, constructs were subcloned into the pET p15A expression vector using NcoI and KpnI restriction sites and the cloning procedure described above. For competitive binding assays, the SR coding sequence at the N-termini of 0 and 10 block entropy ladder constructs was replaced with that of sfCherry^29^ and YFP-OPT2. YFP-OPT2 is a yellow fluorescent protein created by concatenating the sequences of Split YFP 1-10 OPT2^40^ and GFP11 M3 slow^37^. The proteins were cloned using NcoI and KpnI sites, into the pET15A vector, for expression without the N-terminal histidine tag^37^. 1-9 OPT fast and 1-9 M1 slow were subcloned using NdeI and KpnI sites, into the pET p15A vector with an N-terminal histidine tag coding sequence.Transformants were picked and for fluorescent protein expression strains, colonies representing the dominant phenotype were picked and stored for subsequent use, in 25 % glycerol stocks.

For calcium-responsive folding assays, codon optimized gene encoding the intrinsically disordered Repeat in Toxin (RTX) domain, RTX-BR(L) was obtained by gene synthesis (Genewiz, USA) or assembly PCR with overlapping oligonucleotides (Integrated DNA Technologies). These were cloned into a pET28a+ vector using the NcoI and KpnI restriction sites. BL21-DE3 Gold cells were transformed with ligation products.

The integrity of coding sequences for all constructs used in this study were verified by Sanger sequencing.

### *In vitro* complementation assays

Cells were grown at 37 °C, induced with 1 mM B-isopropyl-thiogalactoside-non-animal-origin (IPTG-NAO; VWR USA) in the early-mid log phase ca 1.5 h or OD_600_ of 0.6 and incubated for another 5 h at 37 °C. An Innova 4230 (New Brunswick Scientific) refrigerated orbital shaker incubator was used at 350 rpm for agitation.

Induced cell pellets were lysed in 350 μL of TNG (100 mM Tris-chloride, 150 mM sodium chloride, 10 % glycerol, pH 7.4; all molecular biology grade) buffer by three rounds of sonication in a Biologics 3000 Ultrasonic homogenizer (Biologics, USA) using a 3.1 mm probe. Seven pulses were applied at 50% law and 50% duty cycle, followed by centrifugation at 16100 rcf for 1 min in an Eppendorf 5415D centrifuge (Eppendorf, USA). The supernatant was directly used in assays. Complementation assays of tested constructs were set up as previously described^30^, with the following modifications. Briefly, 40 μL of supernatants were mixed with equal volumes of purified sfCherry^29^ in TNG buffer. 180 μL of refolded GFP1-9 or GFP1-10 was added to 20 μL of the supernatant-sfCherry mixtures. These reactions were set up in 400 μL wells of Nunc Maxisorp white, opaque 96-well flat bottom plates (Nunc, Denmark). GFP1-9, for measurement or GFP1-10, for quantitation was added without a prior ten-fold dilution in TNG buffer, as previously described^30^. qPCR grade adhesive film (VWR, USA) was used to cover and seal all wells. Fluorescence trajectories were recorded using a Biotek Synergy H4 plate reader (Biotek, USA) using the 488/20 nm and 520/20 nm bandpass filters for excitation and emission respectively and a tungsten light source. Sensor gain was set to 100 and 30 measurements were made per well per read. Fluorescence was recorded at 2 min intervals for 12 h. The wells were unilluminated, except for during measurements. sfCherry fluorescence was recorded at the end of the run using 587 nm and 610 nm as the excitation and emission center-wavelengths, respectively, for monochromators. Slit width was set to 9 nm.

Data was collected and processed in Excel spreadsheets. Initial rates of fluorescence evolution were calculated from progress curves with the Excel SLOPE function, using all measurements between 30 and 50 min (10 data points), post-initiation of complementation. The raw fluorescence data for GFP1-9 complementation reactions were scaled for protein concentration data from GFP1-10 complementation reactions. Injection volume variability was accounted for by scaling with sfCherry fluorescence, normalized across the wells.

We fit the data, excluding the 0-chain length construct, to a linear equation:

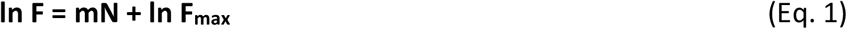

where *N* is the chain length (number of residues), F_max_ is the Y-intercept of the fit and *m* is the parameter for the fit. Using these equations, we calculated *N*_*50*_, the chain length at which:

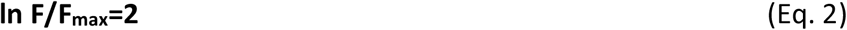

### *In vitro* competitive binding assays

sfCherry and YFP-OPT2 constructs without histidine tags were expressed and extracted using the procedures outlined above for *in vitro* complementation assays. 5 µL of the soluble fractions of cell lysates were run on a 4-20% TGX Mini-Protean gradient gel (Bio-Rad) and stained using Gelcode Blue (Thermo Scientific). Protein abundance was estimated by the intrinsic fluorescence and standardized across samples.

In separate experiments, 400 µL of histidine tagged GFP1-9 M1 at 0.03 µg/µL, purified by inclusion body refolding^30^ was mixed with a solution containing equal volumes (400 µL) of soluble cell lysates containing sfCherry and YFP-OPT2 entropy ladders. In Mix 1, sfCherry was fused to the 0-block (hairpin) entropy ladder and YFP-OPT2 was fused to the 10-block entropy ladder. In Mix 2, the fluorescent proteins were swapped between the two entropy ladder constructs.

After 30 min of incubation, 100 µL of the above mixtures were loaded on 100 µL of TALON resin (Takara Bio, USA), in 200 µL thin-walled PCR tubes and incubated for 5 min at room temperature. Then the resin was pelleted by centrifugation at 800 rcf, followed by supernatant removal. The resin was then washed thrice with 100 µL TNG buffer.

The tubes containing washed resin, were then placed on an Illumatool 9500 epi-fluorescence (Illumatool, USA)^37^ viewing device to image bound protein, using an Olympus C60W digital camera (f/8, ISO 200) (Olympus, Japan). The following filter sets (Table 2) were used to estimate yellow and red fluorescence. The excitation filters were supplied with the Illumatool instrument while the emission filters were made by Edmunds Optics (USA).

**Table 2:** Filter sets used to estimate green, yellow and red fluorescence in competitive binding assays

| Fluorescence filter | Excitation filter (bandwidth) | Emission filter (bandwidth) |
| --- | --- | --- |
| Yellow | 510 nm (10 nm) | 532 nm (10 nm) |
| Red | 550 nm (10 nm) | 610 nm (10 nm) |

**Table 3:** N_50_ values for different combinations of GFP1-9 and S11

| GFP1-9 | S11 | N<sub>50</sub> |
| --- | --- | --- |
| OPT Fast | M4 Fast | 123 |
| OPT Fast | M3 Slow | 51 |
| M1 Slow | M4 Fast | 20 |
| M1 Slow | M3 Slow | 10 |

**Table 4:** Combinations and competing binders used in competitive binding experiments

| Setup | Low entropy species | High entropy species |
| --- | --- | --- |
| 1 | sfCherry-0 block | YFP-OPT2-10 block |
| 2 | YFP-OPT2-0 block | sfCherry-10 block |

### SENP digestion of SUMO-entropy ladder constructs

The *Saccharomyces cerevisiae* SENP protease^38^ was used to digest the SUMO-entropy ladder constructs. The protease was expressed from the pET p15A vector. Equal volumes (100 µL) of cell lysates containing each SUMO-entropy ladder construct and the protease were mixed and allowed to stand for 1 h at room temperature. Negative control samples were diluted with 175 µL of TNG buffer instead of cell lysate containing SENP protease. Digestion was confirmed by running 5 µL of digested or diluted samples on a 4-20% TGX Mini-Protean gradient gel (Bio-Rad), followed by staining with Gelcode Blue (Thermo Scientific) using the manufacturer’s protocol.

### *In vivo* complementation assays

*In vivo* complementation assays were set up between each SR-Entropy Ladder construct and GFP1-9 variant. Complementation with GFP1-10 was used to estimate expression levels of the SR-Entropy ladders by fluorescence^30^.

Overnight cultures were started from the glycerol stocks in a 96-deepwell (2.2 mL/well) plate and grown at 30 °C. Following 14 h of incubation, the cultures were diluted 100,000-fold for plating in zones separated by hydrophobic walls, on a nitrocellulose membrane. The membrane sat atop a selective LB Agar plate. We used sequential induction and imaging procedures described previously^30^, to obtain fluorescence intensities up to 24 h after induction of the detector fragment (GFP1-9 or GFP1-10). Briefly, the colonies were imaged at 30 min intervals on an Illumatool 9500 epi-fluorescence (Illumatool, USA) viewing device, using an Olympus C60W digital camera (f/8, ISO 200) (Olympus, Japan). A 488 nm excitation filter with 10 nm bandwidth and a 515 nm long pass emission filter were used to capture the green fluorescence.

### Calcium-responsive folding assays

Protein expression was performed as described above, for *in vitro* complementation assays. Supernatants from lysates of cells expressing SR-10 block entropy ladder or the RTX-BR(L) protein fusion to the GFP10 and GFP11 tags was diluted 5-fold into 50 mM Tris pH 8.0 and mixed with equal volumes of similarly buffered calcium chloride solutions to final concentrations between 0 and 8 mM using a two-fold dilution series. 1-9 OPT Fast was refolded from inclusion bodies as described above and dialyzed into 50 mM Tris-Cl pH 8.0. Dialysis was performed on 3 mL of the refolded protein in TNG buffer in a 10 kD Molecular Weight Cutoff, Slide-a-Lyzer (Thermo Scientific, USA) cassette. Initially, the protein was allowed to dialyze against 500 mL of buffer at 4 °C for 1 h. Then, the buffer was discarded and a fresh 500 mL of buffer added. Dialysis was continued overnight. The dialyzed protein was similarly mixed with the calcium chloride solutions. The mixed solutions were allowed to stand for at least 5 min to allow protein-calcium equilibration. Then complementation assays were set up in technical triplicates as described above. Samples of sfGFP^41^ in 50 mM Tris pH 8.0 were added to the experiment for instrument drift correction. All complementation reactions were set up on white Nunc Immuno-Maxisorp plates and carried out at room temperature. Complementation rates (V) were calculated as described above and their variation with calcium chloride concentration was fit to a Hill equation^42^, to calculate cooperativity (n) and dissociation constant (K_D_). The following equation was used:

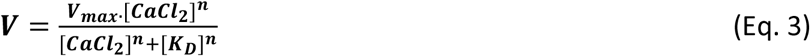

### Peptide and peptoid synthesis reagents

All reagents and solvents deployed were of peptide synthesis or biotech grade. All amino acids were purchased from P3 Biosystems (USA) with the exception of Fmoc His (tBoc), required for high temperature coupling reactions which was obtained from CEM (USA) and Fmoc Norleucine as well as Fmoc Sarcosine which was acquired from Advanced ChemTech (USA). Dimethylformamide (DMF) and the deployed deprotection reagent, 20% Pyrrole (prepared as solution in DMF) was obtained from Alfa Aesar (USA). Pyrrole is the suggested Fmoc deprotection reagent by CEM, as elevated reaction temperatures of 105 °C in deprotections using piperidine lead to loss of side chain protecting groups. The peptide coupling reagents, Diisopropylcarbodiimide (DIC) and Oxyma Pure (Eythylcyanohyroxyiminoacetate) were purchased in peptide synthesis grade from AK Scientific (USA). General reagents such as N,N-diisopropyl ethyl amine (DIPEA), triisopropyl silane ((iPr)_3_SiH; TIPS), thioanisole, octaethylenglycol-dithiol and trifluoroacetic acid (TFA) were purchased from Sigma Aldrich (USA) and methylene chloride (DCM) was obtained from Fisher Scientific (USA). For HPLC purifications, Acetonitrile was acquired from Alfa Aesar (USA) and water was purified in-house (deionized, filtered through a Nanopure to 18.2 MΩ*cm resistivity, and UV-sterilized). Mass spectrometry used highest quality (Optima MS grade) solvents purchased from Fisher Scientific (USA).

### Automated peptomer synthesis using the CEM Liberty Prime microwave peptide synthesizer

A CEM Liberty Prime microwave peptide synthesizer was used for solid phase synthesis at high temperature (105°C). All syntheses were performed at the 0.1 mM scale at the recommended standard instrument chemistry on a Rink amide resin. For short peptides, single coupling instrument cycles were used, with achieved average coupling yields for cycle of > 98.5%. The synthesis deployed 65 s coupling at 105 °C, direct addition of the pyrrole to the hot resin and deprotection at 105 °C for 45 s, 20 s for the three wash steps for a total single couple cycle time of 3 min. Peptoid residues were introduced as preassembled Fmoc peptoid monomers using starting coupling cycle conditions. For longer peptides and peptoids, double coupling of the amino acids was performed and the drain times were increased from 5 to 10 s to accommodate for any resin volume increase over the synthesis cycles. Hydrolysis of acid labile side chain protecting groups during the extended sequence syntheses was prevented by adding 0.1 M DIPEA to the Oxyma solution. High average coupling step yields of microwave assisted syntheses allow even the long and difficult sequences to be obtained in moderate yield.

### Deprotection and removal of the peptomers from the resin

Deprotection used 25 mL of modified “reagent K” mixture: TIPS (1.25 mL/25 mL), thioanisole (0.625 mL/25 mL), octaethyleneglycoldithiol (1.25 mL/25mL), a less odorous substitute for ethylendithiol, and water (1.25 mL/25 mL) in TFA. The resin was pretreated with the quencher solution for 5 min, then TFA was added (to final volume of 25 mL). The deprotections were carried out in 50 mL conical tubes of high-density PP, under a blanket of Argon, for 1.5 h. The solutions were filtered and the filtrate was concentrated to 10 mL. The peptides and peptidomimetics were then precipitated into ice cold ether and collected by centrifugation.

Purification and analysis of synthetic fragments and peptides Purifications (to> 98% +) were performed on a Waters HPLC preparative workstation with 2545 pump (at 20 mL/min) and using a C18 reverse phase column (Waters BEH 130, 5 μm, 19×150) and a gradient from water to acetonitrile with 0.1% TFA from 98% to 20% water/ACN. Peaks were collected based on monitoring at 215 nm using a PDA 2998 detector. Combined product fractions were lyophilized, yielding white fluffy solids. Peptomers were then analyzed for purity by analytical HPLC on a C18 reverse phase column (Waters BEH 130, 5 μm, 4.6×150) with a gradient from 98% to 20 % water-acetonitrile with 0.1% TFA and by mass spectrometry on a Thermo LTQ XL, Thermo Exactive Orbitrap and ABI Sciex 4800 MALDI TOF/TOF mass spectrometers, respectively, in ESI+ mode. Additional data on the mass spectrometry identification are provided in the supplemental (Fig S7-S10).

## Conclusion

We have demonstrated the ability of the tripartite split-GFP system to detect changes in disorder due to changes in the length of a peptide and a moderately hydrophilic peptoid chain. Other similar hydrophilic polymers can therefore be tested using this system. Our experiments with different mutants of two out of three parts of the tripartite system demonstrate the ability to tune the sensitivity of the system. We have also shown the utility to measure environment-dependent changes in the degree of structure in a protein, which can be useful in the development of environmentally responsive polymers, as well as research on cellular signaling that is mediated by similar conformational changes in otherwise intrinsically disordered proteins (IDPs). Lastly, the system can be generally used to discover small molecule regulators of IDPs. These small molecules can be used as drugs and on the corollary, IDP libraries maybe enriched for members that fold around and sequester small molecules, to purify or enrich the latter. The tripartite split-GFP is thus a scalable and direct screening and selection system that can be implemented in protein selection using living cells, like phage and yeast display whilst only requiring translation of the analyte protein. The system is modular, with the small peptides amenable to chemical synthesis. The components can be synthesized at scale and cryo-stored until required. At time of use, site-specific chemistry can be used to label any hydrophilic polymer of interest or a library thereof. Library enrichment is simplified by a single affinity purification of the GFP1-9. We posit that a barcoded library can be enriched *in vitro*, without any living component. Thus this system combines the benefits of a variety of high throughput selection methods for proteins^23–26,43^ and makes them available for selection of abiotic polymers.

## Supplementary Material

**List of items:**

**Table S1: Constructs and their amino acid sequences**

**Table S2: Sequences of constructs used in peptomer experiments**

**Table S3: Analyte and detector combinations used in vitro and in vivo**

**Figure S1: Complementation of RTX-BR(S) and RTX-BR(L) in 50 mM Tris, pH 8.0, with and without 10 % glycerol**

**Figure S2: Titration of calcium against SR-10 block entropy ladder measured with GFP1-9 OPT Fast and GFP11 M4 Fast. Hill binding model fit to the data is shown in dotted black line**.

**Figure S3: Scatter plot showing the complementation of 2 block and 10 block entropy ladder constructs with and without fusion to the SUMO carrier protein**

**Figure S4: SDS-PAGE gel of 2 block and 10 block SUMO-entropy ladder constructs before and after cleavage with SENP/ULP1 protease**

**Figure S5: Live cells after 24 h post complementation**

**Figure S6: Complementation of GFP11 M4 and GFP11-Norleucine with GFP1-10**

**Figures S7-S10: MALDI-TOF spectra of peptomer constructs**

**Table S1:**
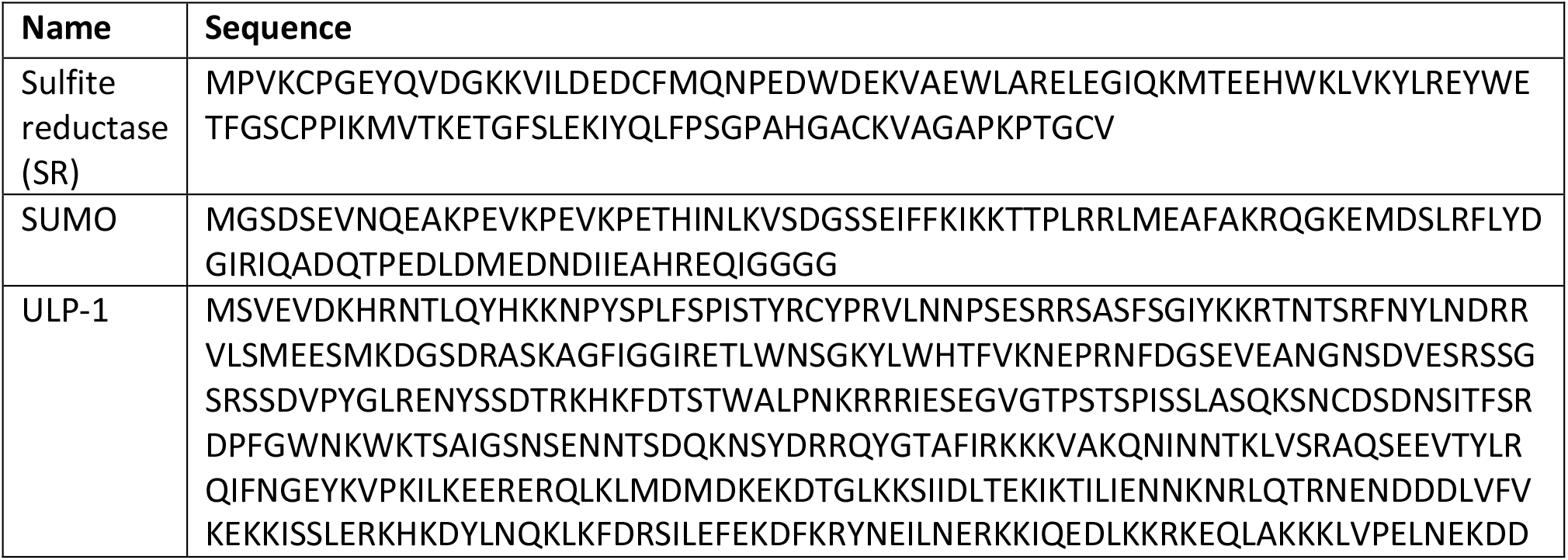
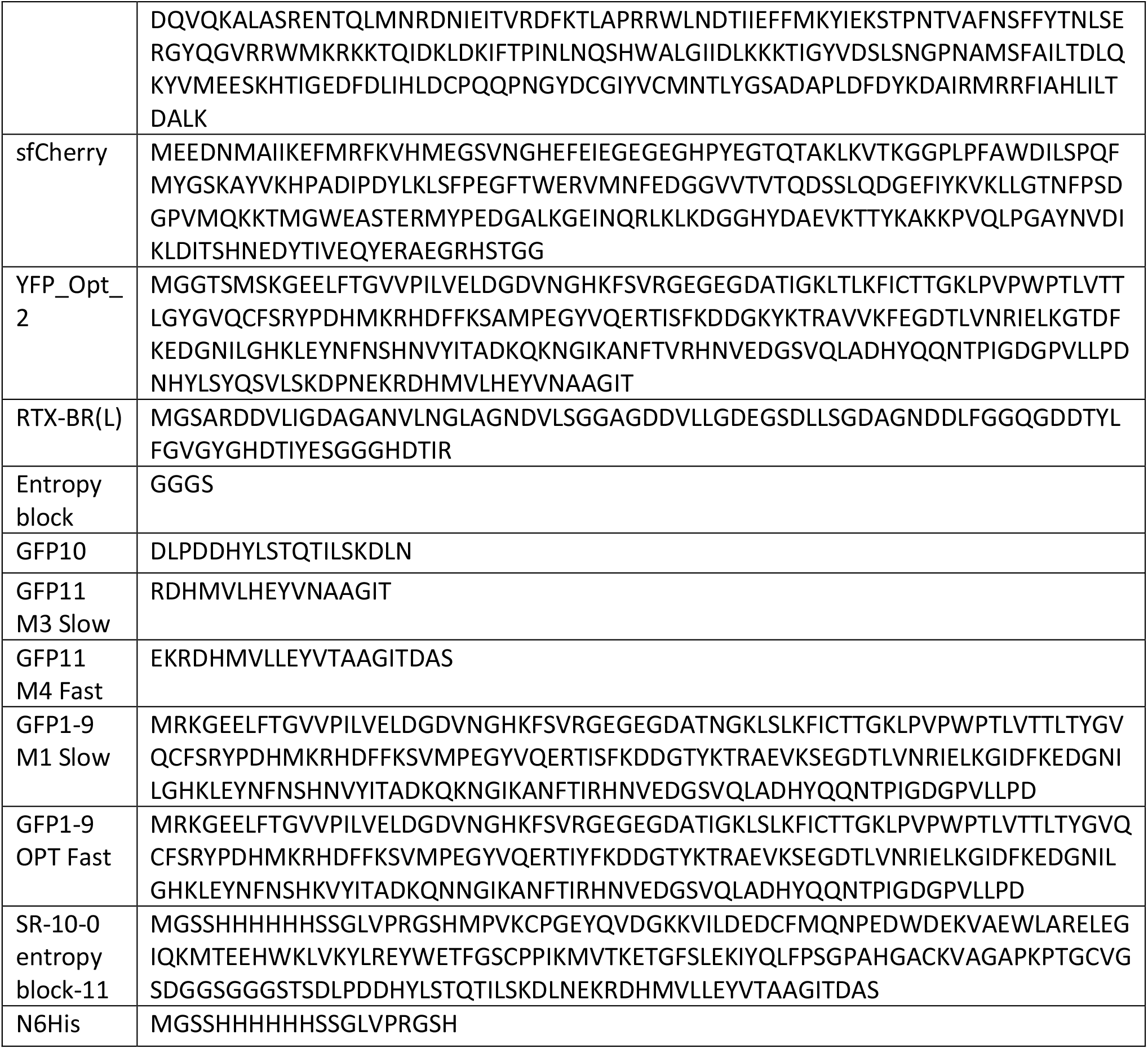
NCBI accession numbers for DNA and amino acid sequences of constructs used in this study

**Table S2:** Sequences of constructs used in peptoid experiments (Sarcosine=Sar, Norleucine=Nlr)

| Construct | Sequence |
| --- | --- |
| Sar3 | DLPDDHYLSTQTILSKDLN-Sar-Sar-Sar- EKRDH-Nrl-VLLEYVTAAGITDAS |
| Sar6 | DLPDDHYLSTQTILSKDLN-Sar-Sar-Sar-Sar-Sar-Sar- EKRDH-Nrl-VLLEYVTAAGITDAS |
| Sar9 | DLPDDHYLSTQTILSKDLN-Sar-Sar-Sar-Sar-Sar-Sar-Sar-Sar-Sar- EKRDH-Nrl-VLLEYVTAAGITDAS |
| Sar12 | DLPDDHYLSTQTILSKDLN-<br>-Sar-Sar-Sar-Sar-Sar-Sar-Sar-Sar-Sar-Sar-Sar-Sar-EKRDH-Nrl-<br>VLLEYVTAAGITDAS |

**Table S3:** Combinations of GFP11, GFP1-9 and GFP1-10 used in this study. The GFP11 was fused to the C-terminus of all analytes

| Analyte | Detector |
| --- | --- |
| 11 M4 fast | 1-9 OPT fast |
| 11 M4 fast | 1-9 M1 slow |
| 11 M3 slow | 1-9 OPT fast |
| 11 M3 slow | 1-9 M1 slow |
| 11 M4 fast | GFP1-10 |
| 11 M3 slow | GFP1-10 |

**Fig. S1:**
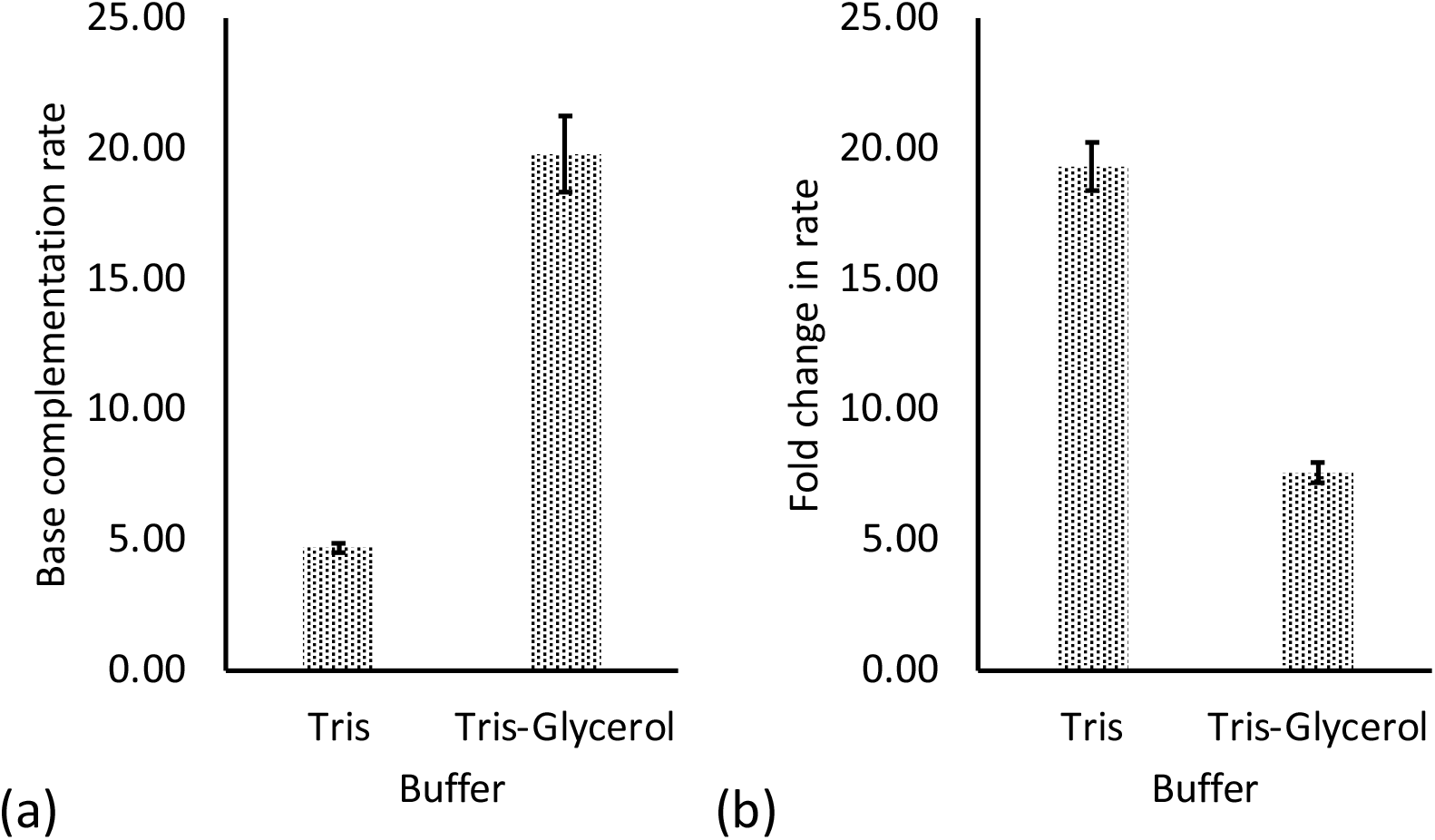
Bar graphs showing the (a) base complementation rate of RTX-BR(S) and RTX-BR(L) in 50 mM Tris, pH 8.0, with and without 10 % glycerol, in the absence of calcium chloride and (b) the fold-change in complementation rate in the presence of 6 mM calcium chloride. Bar heights indicate means of measured values and error bars indicate standard deviation.

**Fig. S2:**
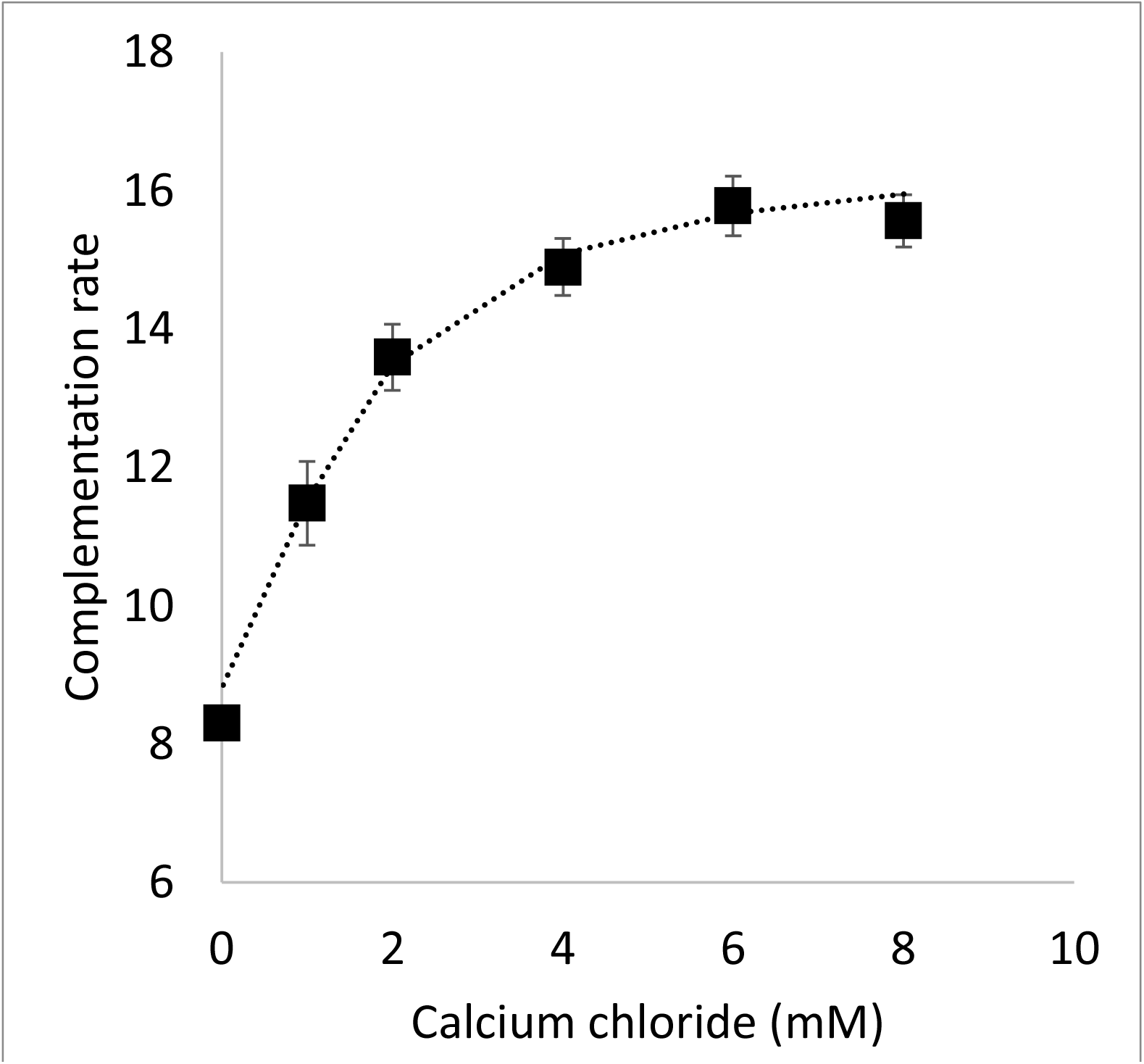
Titration of calcium against SR-10 block entropy ladder measured with GFP1-9 OPT Fast and GFP11 M4 Fast. Points represent means of measurements. Error bars indicated standard deviation. Fit of a Hill binding model to the data is shown in dotted black line.

**Fig. S3:**
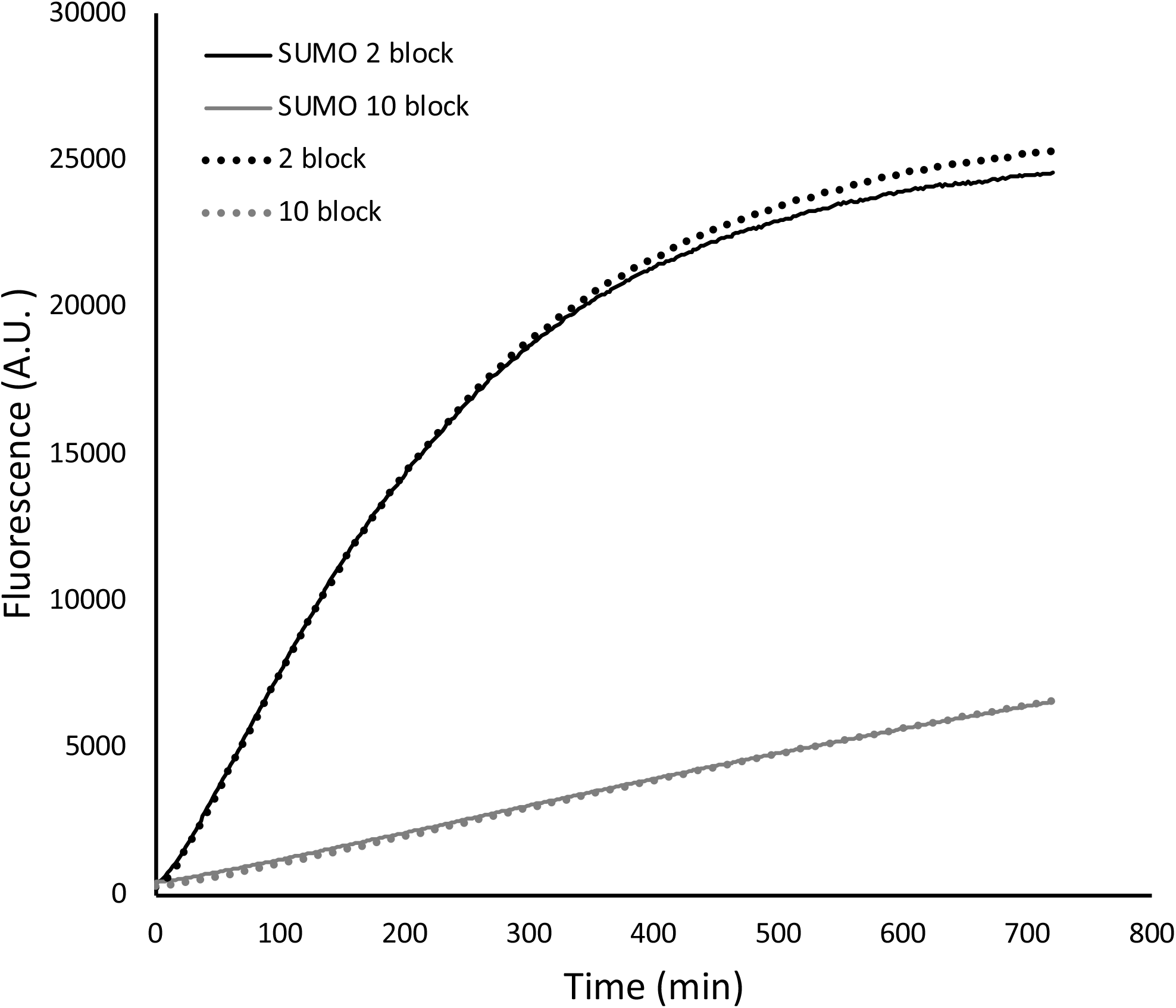
Scatter plot showing the complementation of 2 block and 10 block entropy ladder constructs with and without fusion to the SUMO carrier protein

**Fig. S4:**
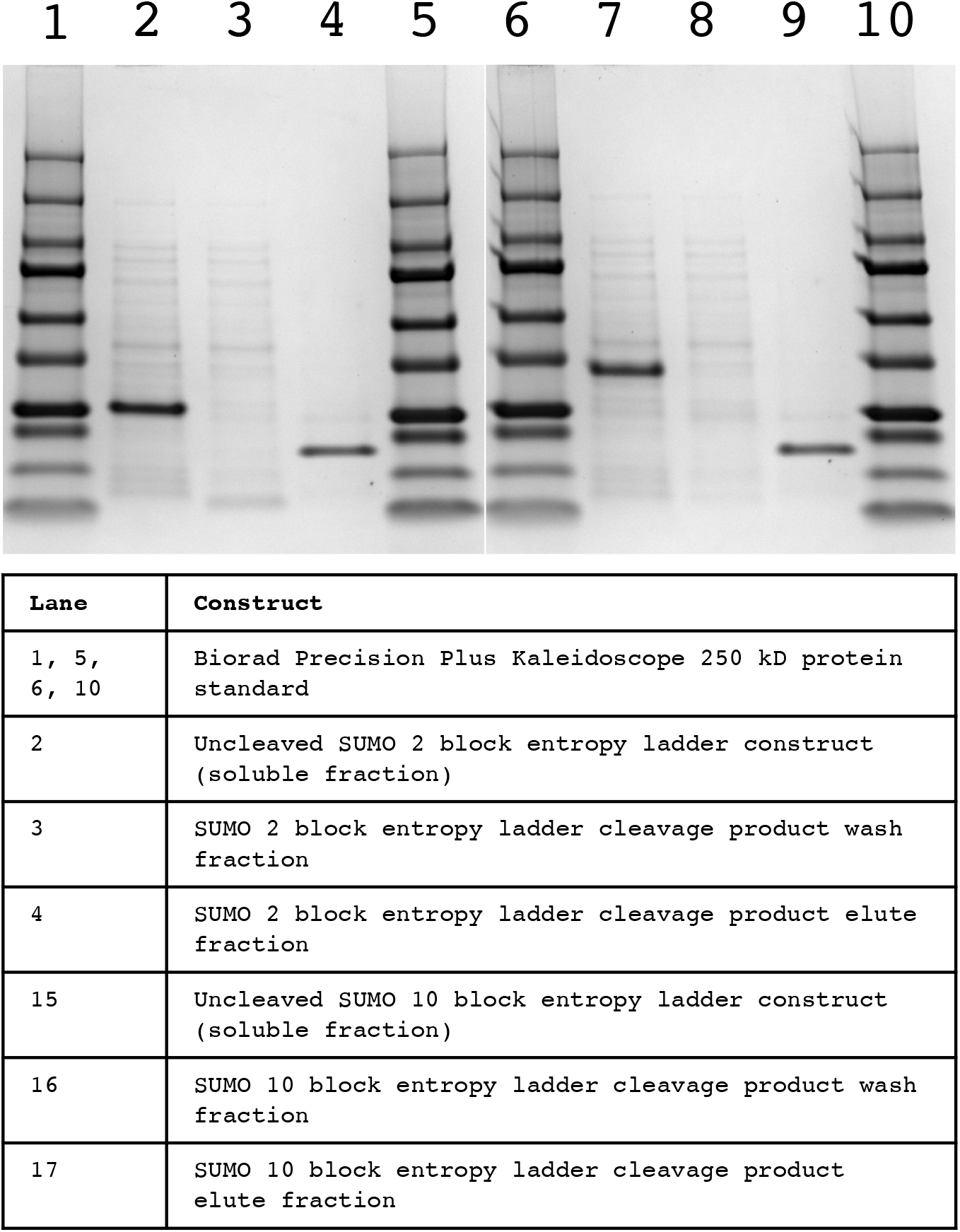
SDS-PAGE gel of 2 block and 10 block SUMO-entropy ladder constructs before and after cleavage with SENP/ULP1 protease

**Fig. S5:**
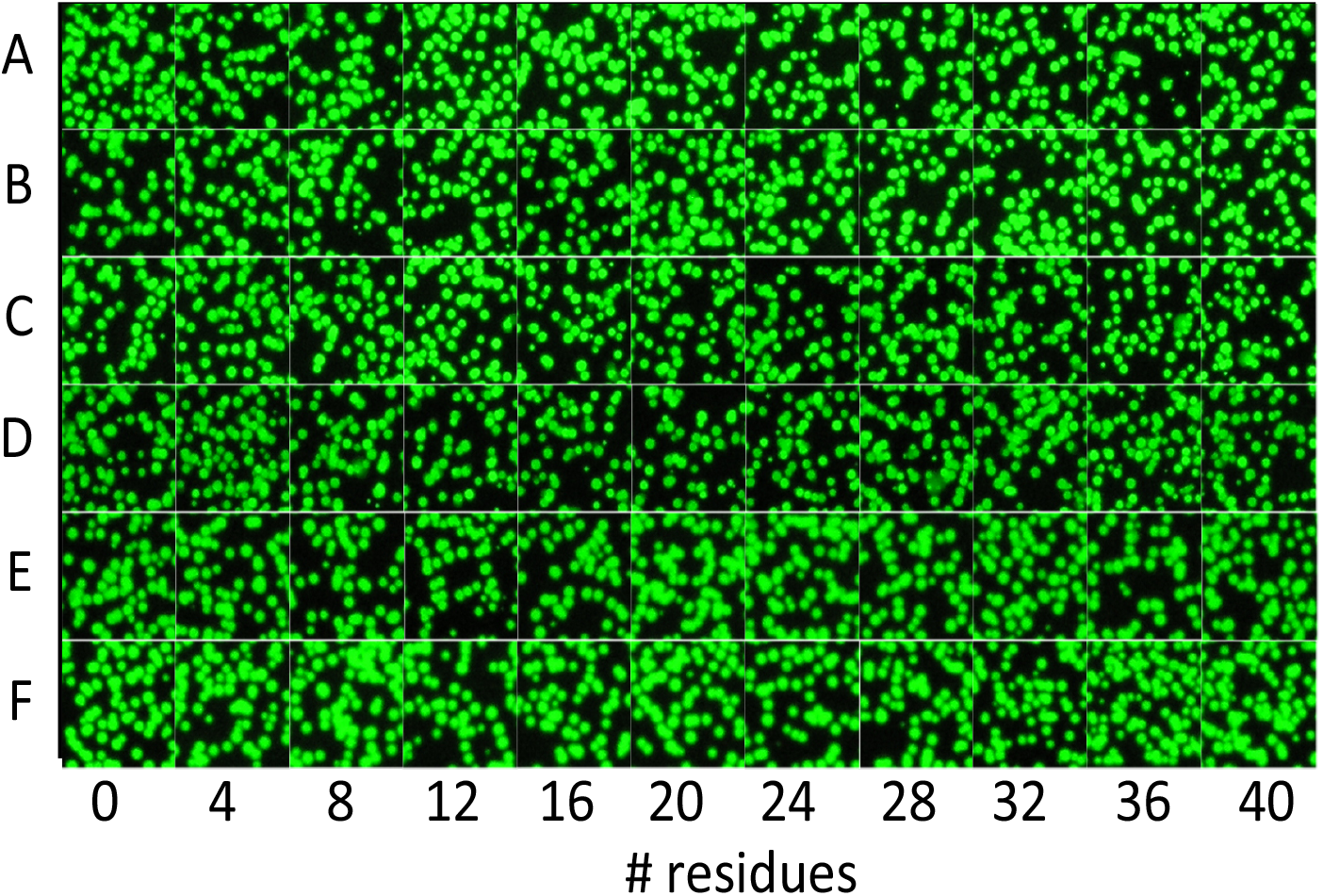
Fluorescent bacterial colonies expressing entropy ladder constructs, after sequential induction of entropy ladder (2 h) and GFP 1-9 or GFP1-10 (24 h). Different rows shown are for cells expressing GFP1-9 or GFP1-10 variants and variants GFP11 at the end of the entropy ladders. (A) 1-9 OPT Fast and 11 M4 Fast, (B) 1-9 OPT Fast and 11 M3 Slow, (C) 1-9 M1 Slow and 11 M4 Fast, (D) 1-9 M1 Slow and 11 M3 Slow, (E) 1-10 and 11 M3 Slow and (F) 1-10 and M4 fast.

**Fig. S6:**
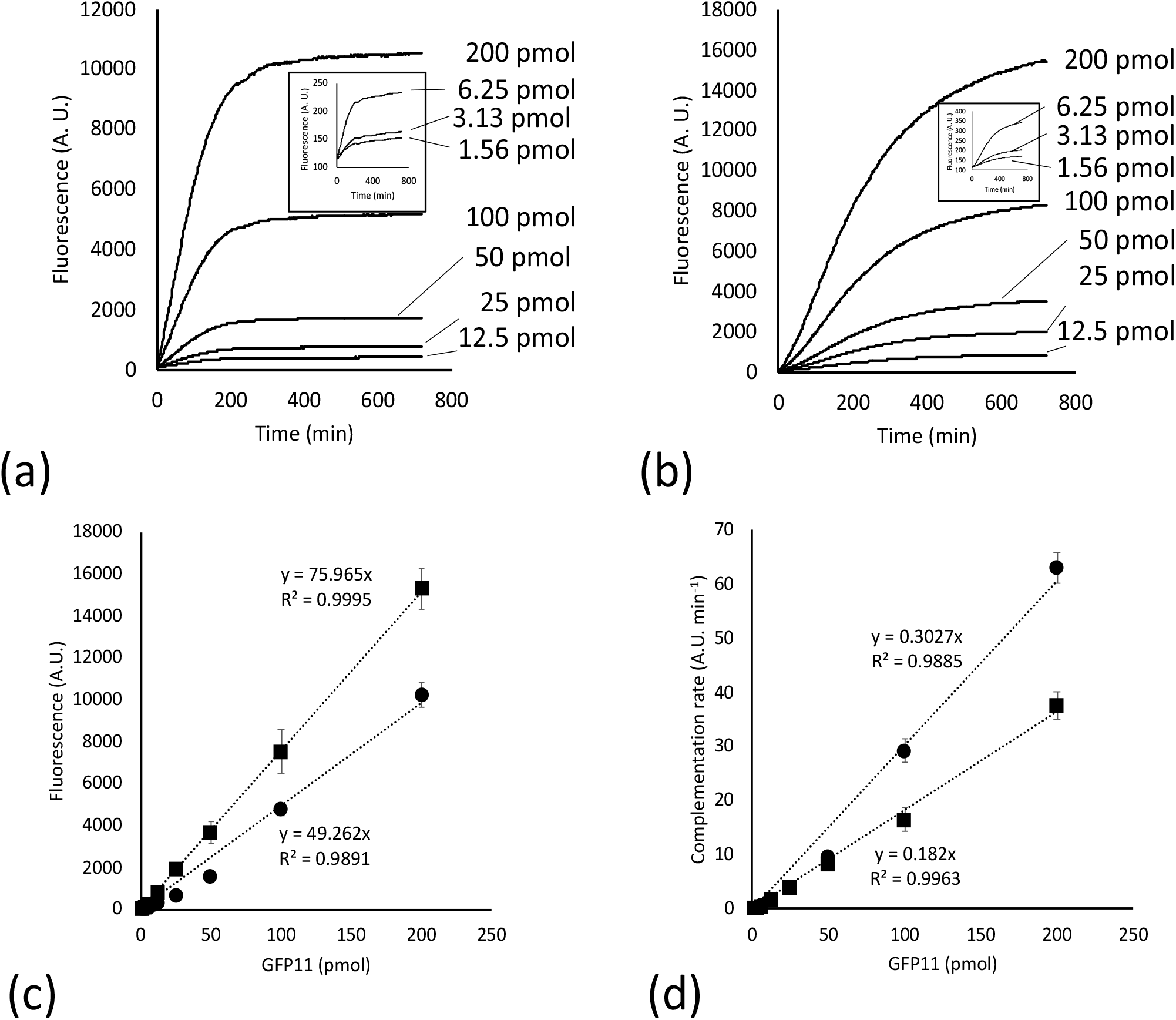
Complementations of GFP1-10 with GFP11 Norleucine and GFP11 M3 at different concentrations. Fluorescence trajectories are shown for GFP11 Norleucine (a) and GFP11 M3 (b). Scatter plots showing the variation of fluorescence after 12 h of complementation (c) and the complementation rate measured between 30 and 50 minutes after onset of complementation (d), with the amount of GFP11.

**Fig. S7:**
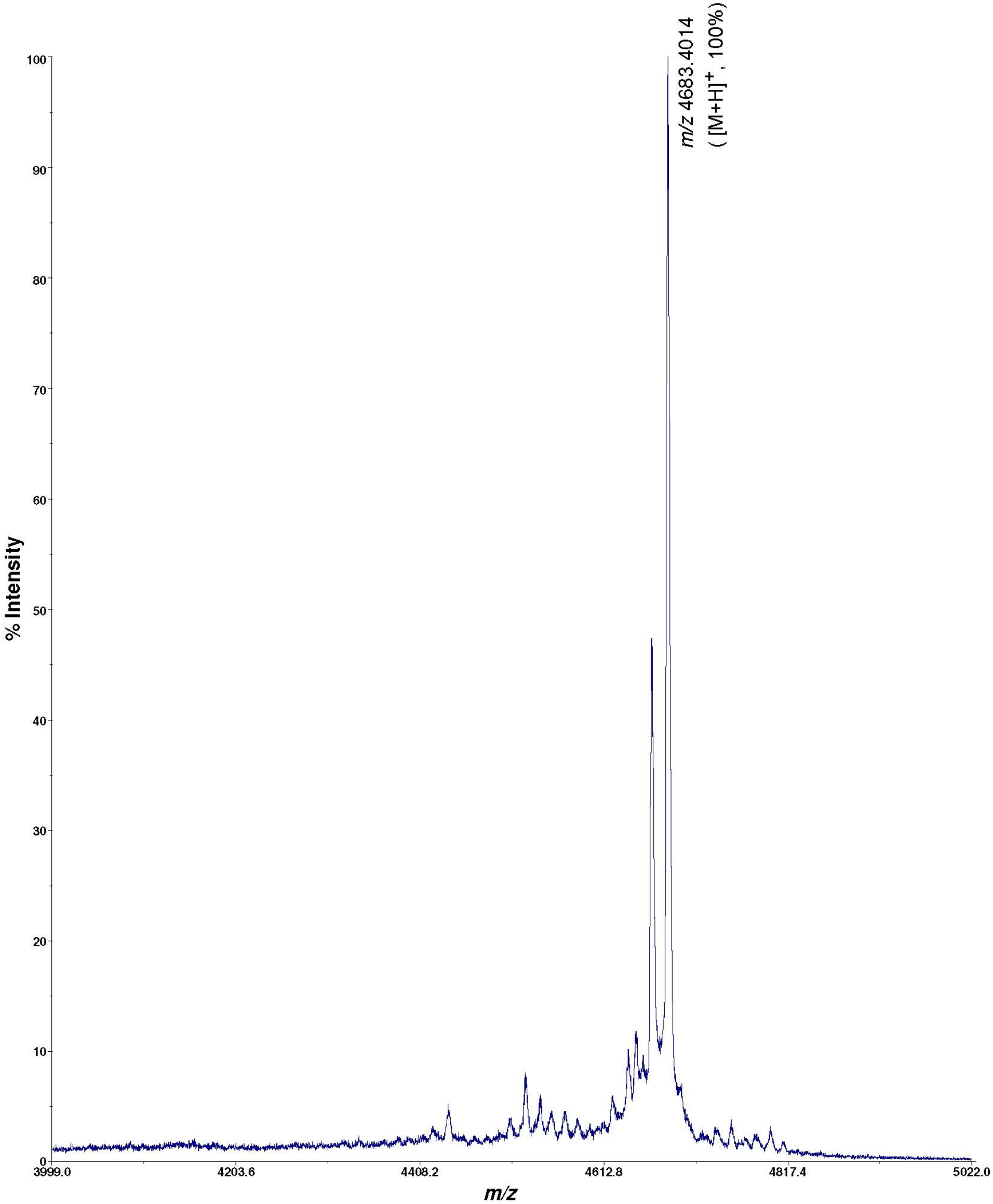
Mass (MALDI) spectrum of 3 block sarcosine peptomer construct showing expected mass to charge ratio of 4683.4

**Fig. S8:**
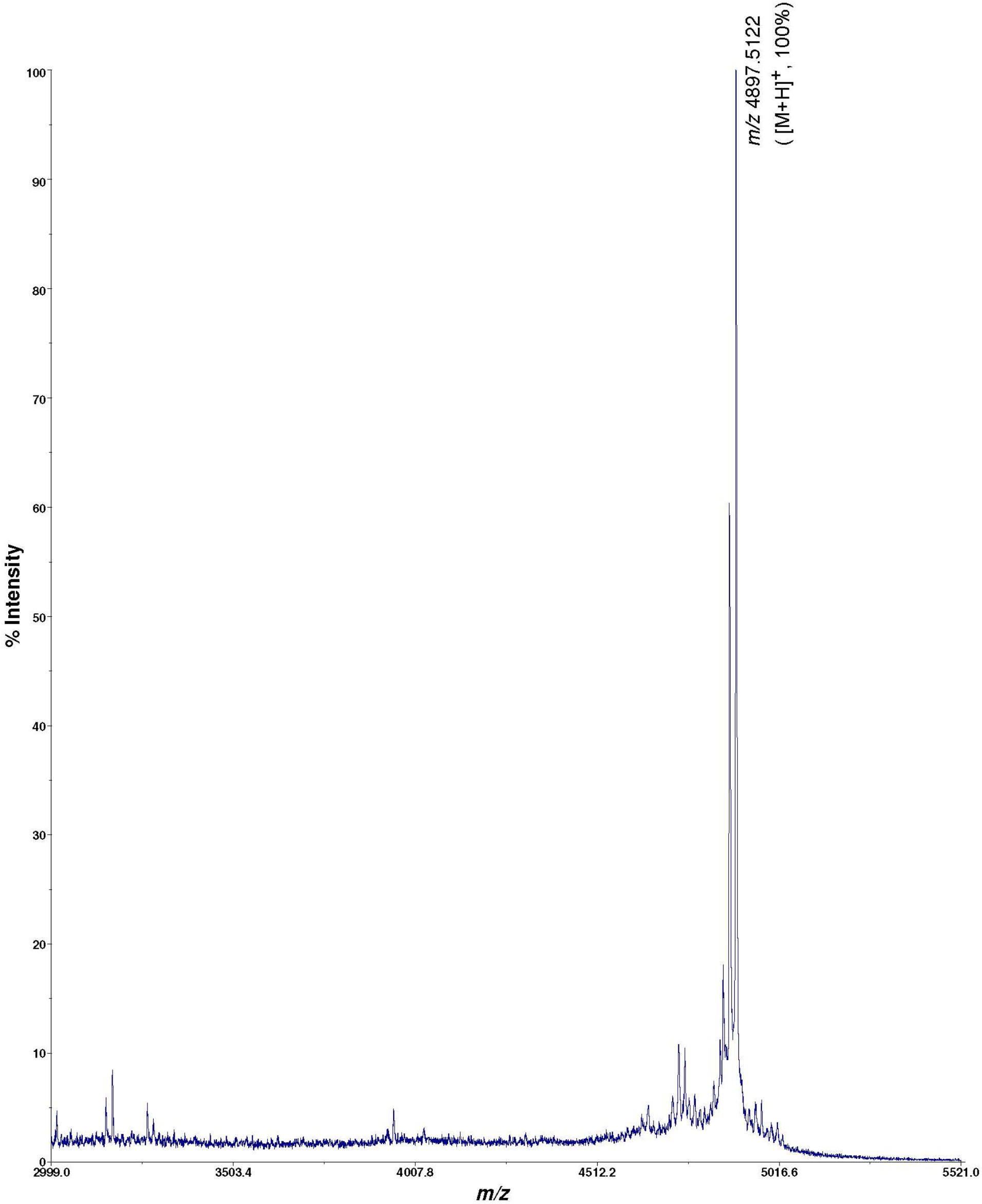
Mass (MALDI) spectrum of 6 block sarcosine peptomer construct showing expected mass to charge ratio of 4897.5

**Fig. S9:**
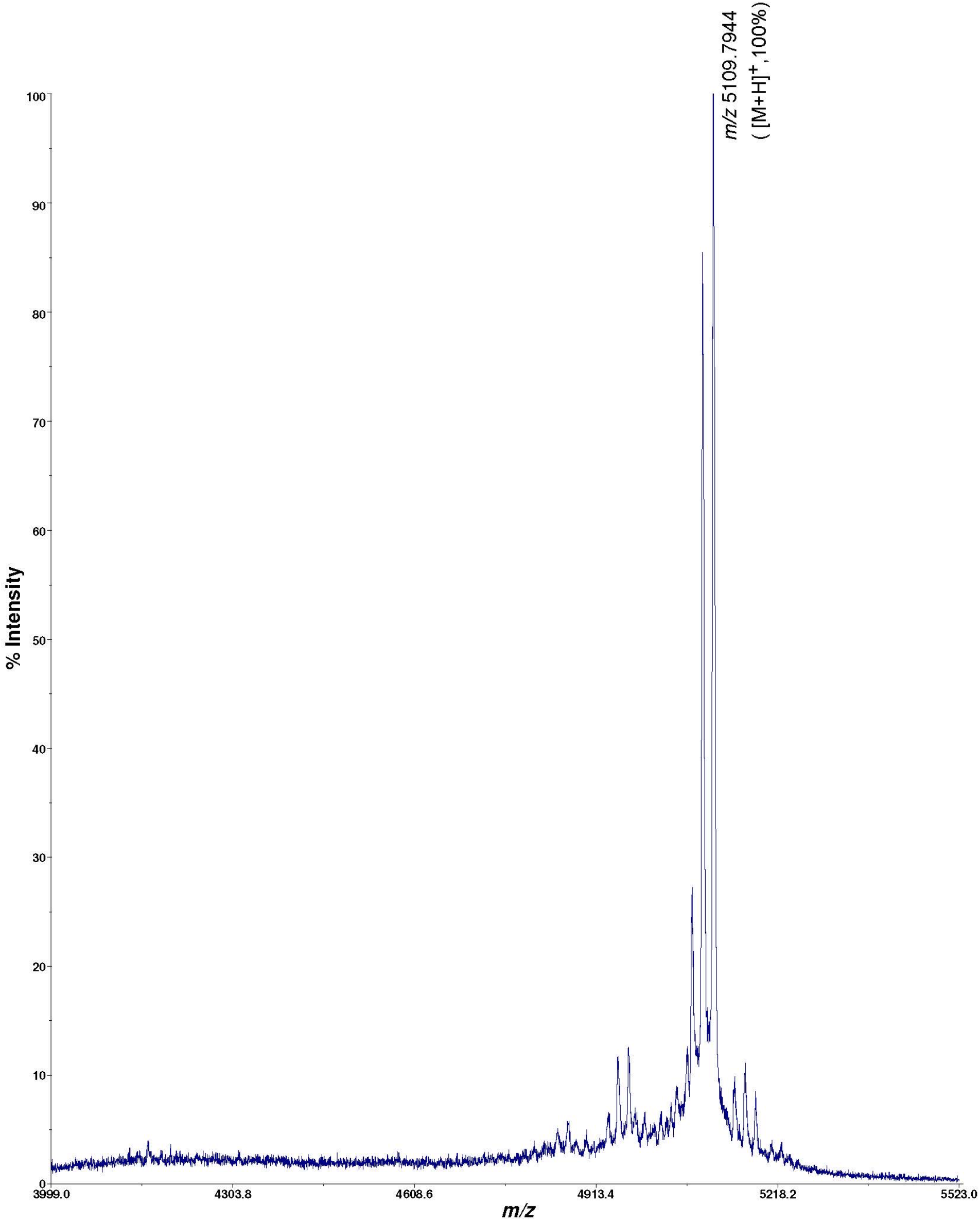
Mass (MALDI) spectrum of 9 block sarcosine peptomer construct showing expected mass to charge ratio of 5109.8

**Fig. S10:**
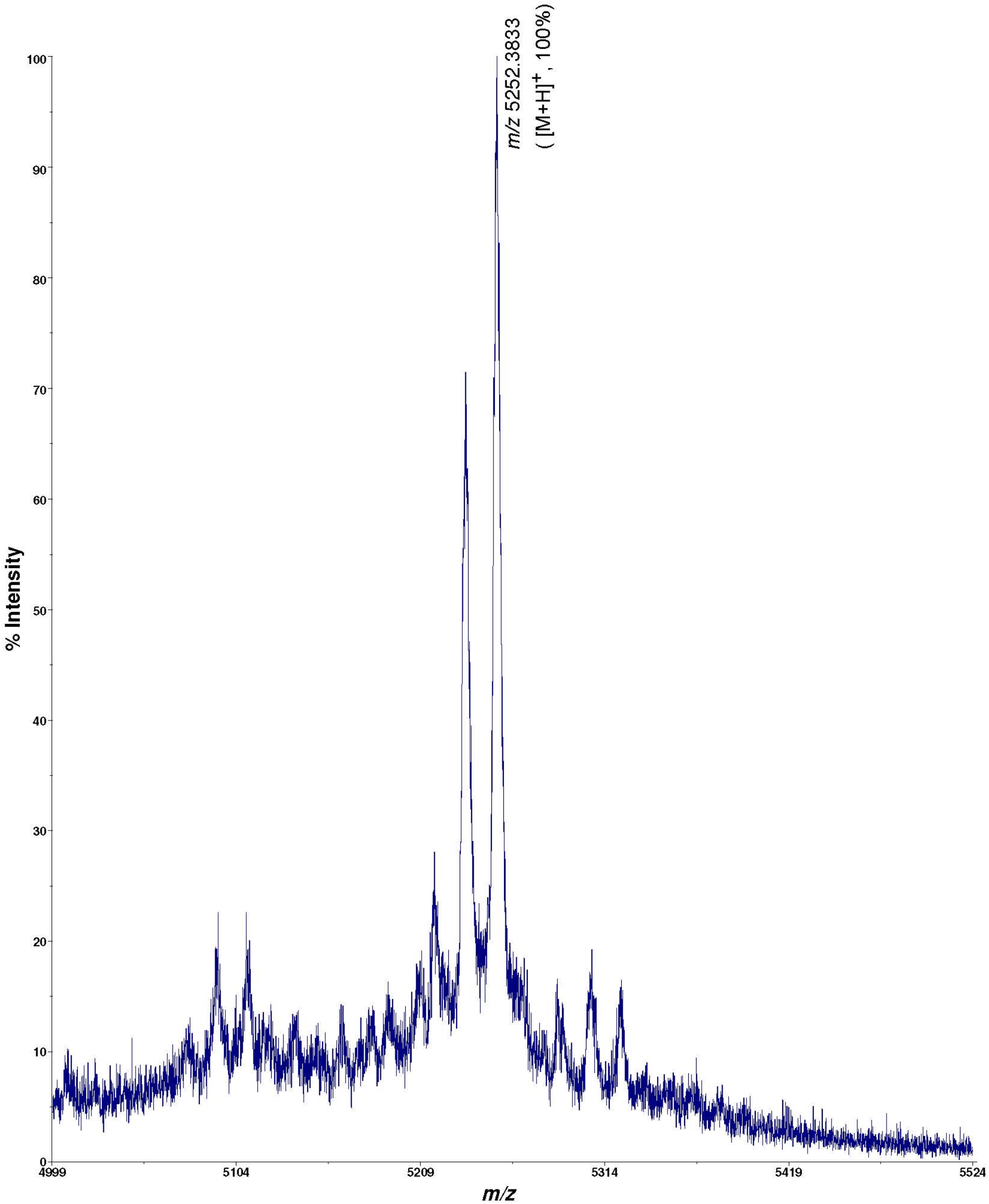
Mass (MALDI) spectrum of 12 block sarcosine peptomer construct showing expected mass to charge ratio of 5252.4

